# Tactile localization of the breast, areola, and nipple

**DOI:** 10.1101/2022.09.14.507974

**Authors:** Katie H. Long, Emily E. Fitzgerald, Ev I. Berger-Wolf, Amani Fawaz, Stacy T. Lindau, Sliman J. Bensmaia, Charles M. Greenspon

**Author notes:** indicates equal contributions (Sliman J. Bensmaia, Charles M. Greenspon).

## Abstract

Touch plays a key role in our perception of our body and shapes our interactions with the world, from the objects we manipulate to the people we touch. While the tactile sensibility of the hand has been extensively characterized, much less is known about touch on other parts of the body. Despite the important role of the breast in lactation as well as in affective and sexual touch, relatively little is known about its sensory properties. To fill this gap, we investigated the ability of women to locate touches on the breast and compared it to that of the hand and back, body regions that span the range of tactile discriminative capabilities. First, we found that the tactile precision of the breast was even lower than that of the back, heretofore the paragon of poor precision. Second, precision was lower for breasts that had undergone greater expansion, consistent with the hypothesis that innervation capacity does not scale with body size. Third, touches to different regions of the nipple were largely indistinguishable, suggesting sparse innervation density. Fourth, localization errors were systematically biased toward the nipple.

**Significance:** Our basic understanding of the tactile capabilities of the breast remains poorly understood in comparison to the hand or face despite the fact that the breast plays a major role in the lives of those with breasts. This paper establishes common methods for studying breast tactile sensation and presents the breast and nipple as two fundamentally discrete tactile units from the torso.

## Introduction

The sense of touch fulfills a variety of different functions in everyday life, from guiding our interactions within the environment to supporting affective communication and sexual function (1). One of the key properties of touch sensations is that they are localized to a specific part of the body: contact on the shoulder produces a sensation experienced on the shoulder, for example. The precision with which we can localize events on the skin has been shown to be determined by the innervation density at that location (2, 3). Because the skin of the fingertips and lips is the most densely innervated, the precision of these body regions is highest, conferring to us an enhanced ability to distinguish touches on the fingers even when they are close to each other. Innervation density and function are intrinsically related: the fingertips are thought to be densely innervated because they account for the vast majority of contacts with objects and precise information about object interactions is critical to dexterous manipulation (4, 5). In contrast, the precision on the back is low because precise localization of a touch on the back is of limited value. Skin surface area also plays a role in innervation density; for example, people with large hands exhibit lower precision than those with small ones (6, 7). This phenomenon is hypothesized to reflect the fact that the number of nerve fibers does not scale with body size: a large body will be more sparsely innervated than a small one given a fixed number of nerve fibers.

While tactile precision has been extensively studied on the limbs and face, precision on the torso has received far less experimental attention (8, 9), with the breast being largely ignored beyond its anatomy (10, 11). To fill this gap, we sought to characterize the spatial precision of the female breast which has roles in lactation, affective touch, and sex. These functions set it apart from other regions of the body. In previous studies, the tactile precision of the breast was found to be comparably low to that of the back and calf (9). However, the experimental approaches have received scrutiny to their susceptibility to inaccuracies (12); only the outer breast was tested (excluding the nipple-areolar complex, NAC), and the relationship with breast size was not assessed, or only examined the NAC but not the outer breast (10). Given that the timeline for breast development extends well past that of nervous system development (13), the female breast offers a powerful test of the fixed innervation hypothesis, which would predict that people with larger breasts would have poorer spatial precision than people with smaller ones.

In the present study, we first measured the tactile precision of two regions of the breast in women– the outer (lateral) breast and medial breast, which includes the NAC and compared these to the hand and back. Second, we examined the relationship between the precision of the outer breast and breast size. Third, we examined women’s ability to judge the absolute location of touches to their breast. We first found that the spatial precision of the breast is very low, with similar sensitivity to the back. Second, spatial precision is inversely correlated with breast size, as predicted from the fixed innervation hypothesis. Third, touches to different parts of the nipple are indistinguishable, suggesting sparse tactile innervation. Fourth, touches on the outer breast are systematically mis-localized as being closer to the nipple than they actually are.

## Results

### The breast has low spatial precision

We first measured each participant’s ability to judge the relative position of two touches applied in succession at two nearby locations on the skin of the hand, back, and breast using a punctate probe. The first of the two touches (the reference) was at the same position on each trial, and the second touch (the comparison) was either above or below the first at a pre-specified distance. For the hand, distances ranged from 1 to 10 mm; for the other body regions, distances ranged from 2.5 to 40 mm, anticipating lower precision based on preliminary testing (**Figure 1**). The participant’s task was to report whether the test stimulus was located above or below the reference stimulus. We then assessed the participant’s performance as a function of the distance between the test and reference and characterized the distance from the reference required to reliably locate the test stimulus – the just noticeable difference (JND) – for each participant and region.

**Figure 1.**
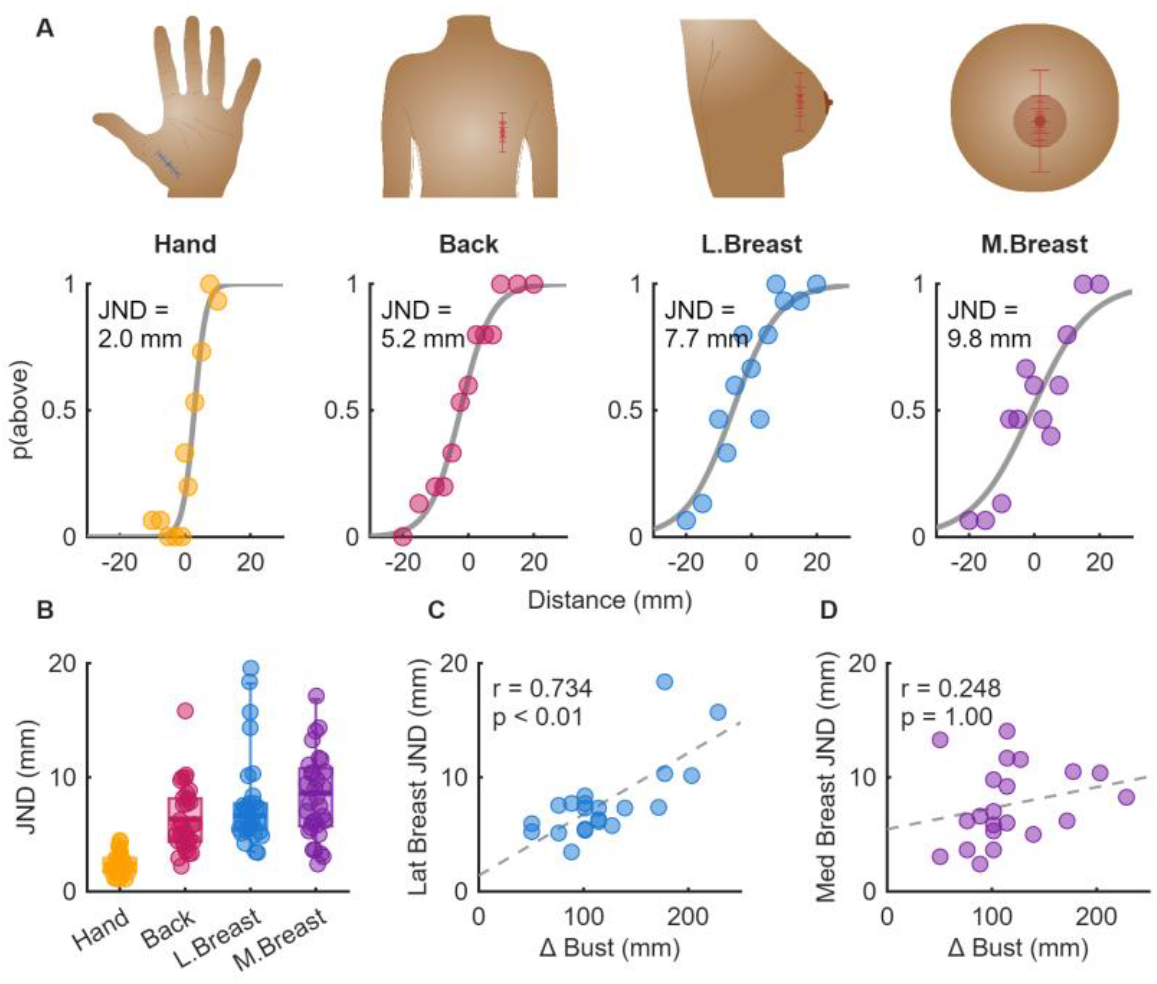
Location discrimination. **A**| Example psychometric functions for one subject on the location discrimination task for the hand, back, outer breast, and medial breast (including NAC). Negative values denote test points “below” the reference and the JND indicates the distance at which the subject could reliably (75% trials) locate the stimulus. **B**| Distribution of JNDs for each subject at each region. **C**| Relationship between the difference in size between the bust and underbust (Δ Bust) and the JND of the lateral breast and **D**| medial breast, respectively.

As expected, the hand demonstrated significantly greater spatial precision (median [25^th^,75^th^] percentiles: 2.16 [1.73, 2.93] mm) than either the back (5.72 [4.36, 8.16] mm) or either breast region (6.63 [5.41, 7.73] mm and 8.00 [5.72, 10.80] mm, for the lateral and medial breast respectively) as indicated by the lower JNDs (**Figure 1A, B**, 1-way ANOVA: F_[3,126]_ = 22.7, p < 0.01; Tukey’s HSD post-hoc t-test: p < 0.01). Interestingly, the medial breast yielded even shallower functions and lower spatial precision than did the back (**Figure 1A, B**, 1-way ANOVA with Tukey’s HSD post-hoc t-test: p = 0.0284), though no statistical differences were observed between the lateral breast and either the back or medial breast (p = 0.3974 and 0.7377, respectively). In other words, touches needed to be between 3 and 4 times as far apart on the breast than on the hand to yield equivalent location discrimination performance.

### Tactile precision is worse for women with larger breasts

The spatial precision of the hand has been shown to depend on the size of the hand, with smaller hands yielding better precision (6). With this observation in mind, we investigated whether the inter-participant differences in breast precision might be driven in part by differences in breast size. To determine the relative expansion (increase in surface area) of each participant’s breast, we computed the difference between their bust (circumference of the torso at the nipple line) and underbust (circumference of the torso at the inframammary fold). Comparing this value with their spatial precision, we found that the precision of the lateral breast decreased (JNDs increased) as breast size increased (**Figure 1C**, Pearson’s correlation with Bonferroni PHC (n=4): r = 0.734, p < 0.01), a phenomenon that did not extend to the medial breast (**Figure 1C**), hand, or back (**Supplementary Figure 1**, p > 0.05 for all), implying that any expansion is preferentially limited to the skin of the lateral breast and that the observed relationship between breast size and precision was not spurious.

### Tactile events are poorly discriminated on the nipple

While the result that the medial breast has the lowest spatial resolution is not necessarily surprising given the lack of mechanoreceptors near the surface of the skin (14), the magnitude of the correlation was unexpected. Indeed, we observed that the JNDs for each participant were, on average, only marginally smaller than the measured diameter of the nipple (80% ± 52%). This would imply that 2 points on opposite halves of the nipple (80% diameter apart) might be confused as the same point. Given that the NAC is inherently a heterogeneous structure as it is composed of three anatomically distinct regions of the breast (the nipple, the areola, and the nearby outer breast), we reasoned that this approach may be biased and thus not reflect the functional ability to locate sensations on the breast.

To assess if participants could reliably perceive tactile events at the level of either the areola or nipple, we delivered punctate touches to each quadrant on the nipple or areola and the participant reported the quadrant in which the touch had been delivered (**Figure 2A**). Specifically, we marked the edge of the nipple or the 50% point between the center of the nipple and the edge of the areola in each of the cardinal directions such that the proportional distance was constant across participants. When touching different quadrants of the areola, participants were able to reliably locate stimuli (**Figure 2B**, mean ± standard deviation: 55% ± 21% where chance is 25%, one-sided Monte-Carlo test, Z = 15, p < 0.01), with 8 of the 10 participants performing significantly above chance (p < 0.05 after Bonferroni PHC where n = 10). Of the remaining participants, one performed marginally above chance (40%, p = 0.25 after Bonferroni PHC) while the other systematically misperceived the location and performed well below chance (7.5%, p = 1 after Bonferroni PHC). On the nipple, however, participants were consistently worse at locating stimuli than the areola (Wilcoxon signed-rank test, p = 0.0137) where only 3 of the 10 participants outperformed chance, though the group as a whole outperformed chance (**Figure 2B**, 36% ± 13%; Z = 5.5, p < 0.01). This finding suggests that the ability to locate tactile events is significantly worse at the nipple than at the areola, consistent with equivalent innervation and the smaller size of the nipple, and re-affirms our earlier finding that spatial precision at the medial breast is poor. Finally, to determine if this task was also influenced by spatial expansion, we sought to compare the breast size with performance on the quadrant task. Unfortunately, only 4 of the 10 participants returned for measurement and while negative trends were observed, the sample size is insufficient for formal analysis.

**Figure 2.**
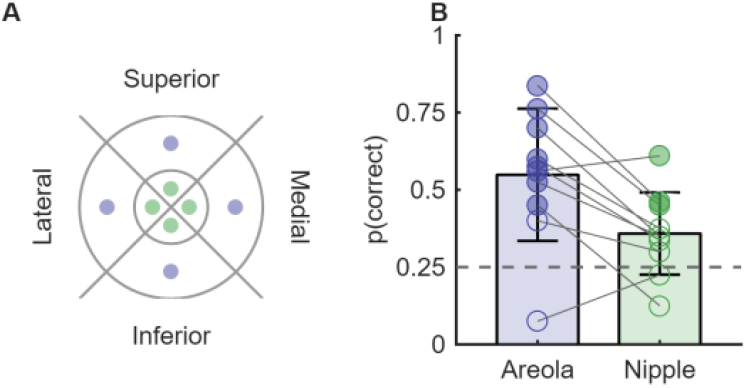
Spatial discrimination at the nipple and areola. **A**| Test locations for the quadrant discrimination task. The outer ring represents the areola and the inner ring the nipple. The subject reported location using a number from 1 to 4 progressing clockwise from “above.” **B**| Proportion correct for participant at each test location. Solid markers indicate participants whose performance was significantly above chance (N=10).

### Localization of tactile events on the breast is biased towards the nipple

In the experiments described above, we investigated the participants’ ability to distinguish the relative locations of two touches to the breast. Next, we examined their ability to identify the absolute location of a tactile stimulus on their torso. To this end, we presented a single touch to the breast or the back with a punctate probe at one of 25 locations after which each participant marked the location of the touch on a three-dimensional digital image of the participant’s own breast or back (**Figure 3A, B**).

**Figure 3.**
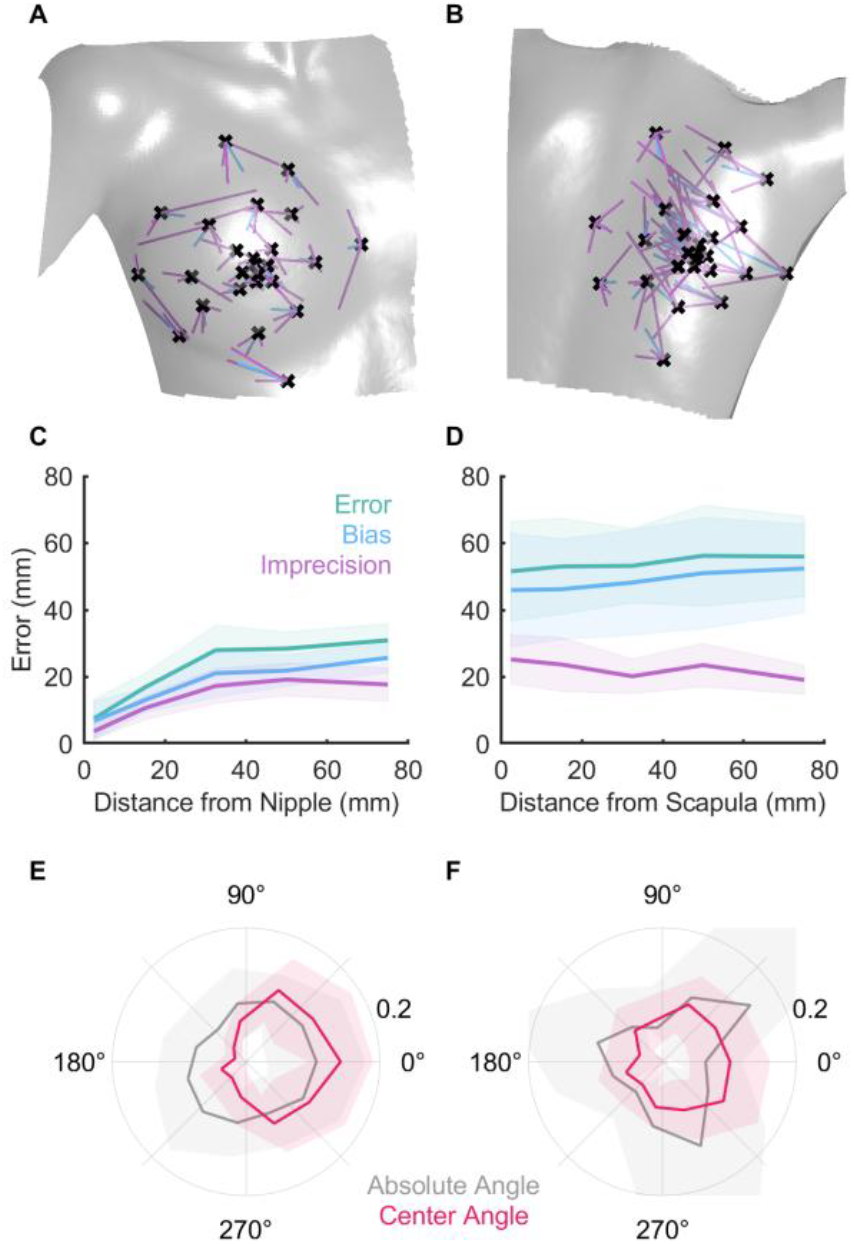
Absolute localization of contact events on the breast and back. **A**| Example localization task data for the breast or **B**| back of one participant. Gray surface represents scanned torso. Black crosses indicate the true location of each stimulus, purple lines indicate the vector between the stimulus location and reported location, and blue lines indicate the vector between the stimulus location and average reported location across blocks. **C**| Reporting error (3D Euclidean distance) as a function of distance across participants for the breast and **D**| back. Error indicates mean error for each trial (mean of purple vector length in **A**,**B**), bias indicates the error of the average response (blue vectors in **A**,**B**), and imprecision is the mean pairwise error between individual responses for each point (distance between the end of purple vectors in **A**,**B**). **E**| The distribution of angles (2D, no depth axis) between the stimulus location and the reported location (gray) or the difference between said angle and the angle towards the relevant landmark (red) for the breast (nipple) and **F**| back (scapula).

To estimate the degree to which participants could accurately localize touch events, we computed the error for each body part across participants and found that, contrary to the precision task, participants had lower errors on their breasts in comparison to their back (**Supplementary Figure 2A;** paired t-test, t[9] = 7.4, p < 0.001). Next, we computed this error as a function of the distance from the center landmark for each region (nipple or the inferior angle of the scapula, i.e. the bottom corner, **Figure 3C, D**) and found that in the case of the breast, the errors increased systematically as the distance from the nipple increased consistent with the quadrant localization task (Linear Mixed Model, LMM: distance: n = 248, B = 0.234, t = 9.194, p < 0.001, **Table 1**). This association was not found for the back (distance: n = 248, B = 0.093, t = 1.714, p = 0.087, **Table 2**). Then, using an LMM to compare the groups, we found both a significant interaction between the effect of distance and the location (location x distance: n = 496, B = -0.139, t = -2.723, p = 0.006, **Table 3**). Given that any errors are likely a result of two sources of noise: bias and response variability (imprecision), we next sought to quantify the relative contribution of the two sources. We estimated the bias for each point by computing the centroid of all reports for a given stimulus across trials and measured the error between the centroid and the stimulus location. To measure the imprecision, we computed the mean pairwise distance between each of the reported locations for a given stimulus location and the mean of those reported locations (**Figure 3C, D**). We found that, in both the breast and back, the error that originated from systematic bias significantly outweighed that of the imprecision (location x error type: n = 995, B = -23.05, t = 12.728, p < 0.001).

**Table 1:**
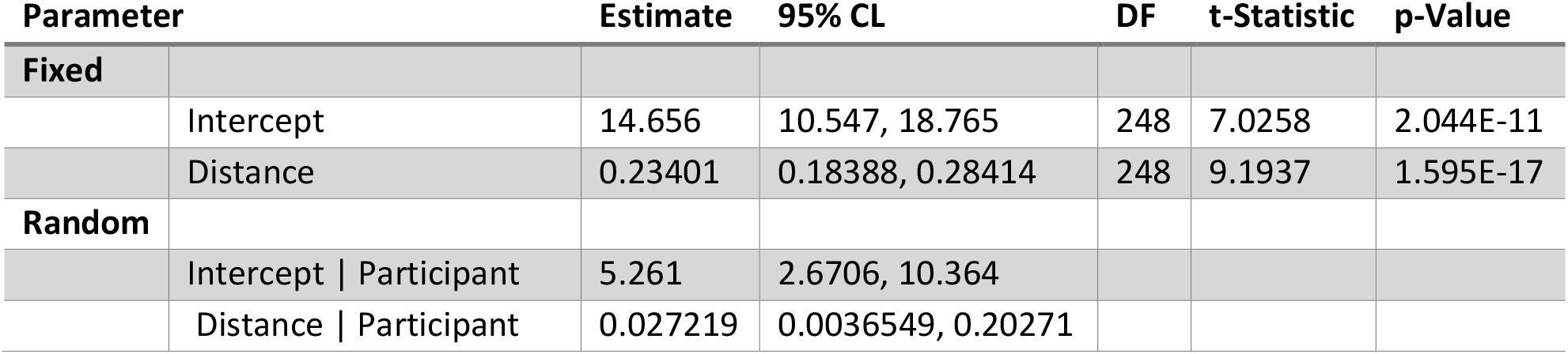
Linear mixed model for the relationship between distance and error for tactile localizations on the breast. *error = distance + (distance* | *participant)*

| Parameter |  | Estimate | 95% CL | DF | t-Statistic | p-Value |
| --- | --- | --- | --- | --- | --- | --- |
| **Fixed** |  |  |  |  |  |  |
|  | Intercept | 14.656 | 10.547, 18.765 | 248 | 7.0258 | 2.044E-11 |
|  | Distance | 0.23401 | 0.18388, 0.28414 | 248 | 9.1937 | 1.595E-17 |
| **Random** |  |  |  |  |  |  |
|  | Intercept \| Participant | 5.261 | 2.6706, 10.364 |  |  |  |
|  | Distance \| Participant | 0.027219 | 0.0036549, 0.20271 |  |  |  |

**Table 2:** Linear mixed model for the relationship between distance and error for tactile localizations on the back. *error = distance + (distance* | *participant)*

| Parameter |  | Estimate | 95% CL | DF | t-Statistic | p-Value |
| --- | --- | --- | --- | --- | --- | --- |
| **Fixed** |  |  |  |  |  |  |
|  | Intercept | 49.668 | 41.571, 57.764 | 248 | 12.083 | 9.7547e-27 |
|  | Distance | 0.093385 | -0.013932, 0.2007 | 248 | 1.7139 | 0.087801 |
| **Random** |  |  |  |  |  |  |
|  | Intercept \| Participant | 10.934 | 5.9341, 20.145 |  |  |  |
|  | Distance \| Participant | 0.11002 | 0.036619, 0.33057 |  |  |  |

**Table 3:** Linear mixed model for the relationship between distance and error for tactile localizations on the breast and back. *error = distance + location + (distance x location) + (distance* | *participant)*

| Parameter | Estimate | 95% CL | DF | t-Statistic | p-Value |
| --- | --- | --- | --- | --- | --- |
| **Fixed** |  |  |  |  |  |
| Intercept | 14.601 | 8.3922, 20.809 | 496 | 4.6206 | 4.8848e-06 |
| Distance | 0.23491 | 0.13881, 0.33101 | 496 | 4.8027 | 2.0762e-06 |
| Location | 34.927 | 29.63, 40.225 | 496 | 12.954 | 2.8993e-33 |
| Distance x Location | -0.13876 | -0.23889, -0.038634 | 496 | -2.7229 | 0.0066996 |
| **Random** |  |  |  |  |  |
| Intercept \| Participant | 7.9445 | 4.5113, 13.99 |  |  |  |
| Distance \| Participant | 0.10285 | 0.049394, 0.21416 |  |  |  |

**Table 4:** Linear mixed model for the relationship between distance and error for tactile localizations on the breast and back with respect to error type (bias or imprecision). *error = distance + location + error type + (error type x location) + (distance* | *participant)*

| Parameter | Estimate | 95% CL | DF | t-Statistic | p-Value |
| --- | --- | --- | --- | --- | --- |
| **Fixed** |  |  |  |  |  |
| Intercept | 15.291 | 11.301, 19.281 | 995 | 7.5203 | 1.2187e-13 |
| Distance | 0.10133 | 0.046744, 0.15592 | 995 | 3.6427 | 0.00028369 |
| Location | 29.034 | 26.52,31.548 | 995 | 22.663 | 5.1594e-92 |
| Error Type | -4.0629 | -6.5756,-1.5503 | 995 | -3.1731 | 0.0015544 |
| Location x Error Type | -23.048 | -26.601, -19.495 | 995 | -12.728 | 1.7171e-34 |
| **Random** |  |  |  |  |  |
| Intercept \| Participant | 5.2017 | 2.942, 9.1969 |  |  |  |
| Distance \| Participant | 0.069106 | 0.033314, 0.14335 |  |  |  |

As bias was prevalent in the localization task, we next sought to determine if the bias was systematic across stimulus locations. Consequently, we assessed the tendency for the error vector to be uniformly distributed in two dimensions (**Supplementary Figure 2B-C**). Importantly, because the back is a relatively flat surface in comparison to the breast, error vectors were only computed in the horizontal plane and depth was excluded when computing the angular error. When computing the absolute angle between stimulus location and the average reported location (blue vectors in **Figure 3A, B**) and computed the distribution across participants (**Figure 3E, F**). Across participants, the distribution of biases did not significantly deviate from uniformity (permutation test for uniformity: R = 0.1314, p = 0.9331 and R = 0.1856, p = 0.7171, respectively). Next, when we assessed if the bias with respect to the center point (nipple or scapula) was consistent, we found that both the breast and back tended to exhibit biases (R = 0.3140, p = 0.0100 and R = 0.6164, p < 0.0001, respectively), with the effect roughly doubled for the breast.

To further quantify this observation, we computed the vector strength – a measure of circular uniformity-of the biases for individual participants. For the breast we did not detect significant non-uniform distributions across participants, though some participants demonstrated biases (e.g. towards the top of the shoulder, **Supplementary Figure 2D-F**; vector strength Monte-Carlo test: p < 0.05 for 4/10 participants). Examining the back, significant non-uniform distributions were observed for all but 1 participant, however, the direction was inconsistent across participants implying that reports were not biased towards any single landmark. Next, we computed the distribution of relative angles between the bias vector and the center point of our task and found that 7 of 10 participants were biased towards the nipple (vector strength Monte-Carlo: p < 0.05) and 2 were biased towards the scapula, suggesting that while individual biases are present, they vary substantially across the population.

Given these findings, we conclude that the breast has lower tactile precision than the hand and is instead comparable to the back. Moreover, localization of tactile events to both the back and breast are inaccurate but localizations to the breast are consistently biased towards the nipple.

## Discussion

First, we found that the spatial precision of the lateral breast – excluding the nipple and areola – is almost four times lower than that of the hand, and even lower than that of the back, previously considered the epitome of poor precision. Second, the nipple has such low tactile precision that touches to different aspects of the nipple are nearly indistinguishable from one another. Third, the precision of the breast tends to be poorer for women with large breasts, consistent with the theory that innervation capacity is fixed. Together, the data indicates that the nipple constitutes a landmark on the breast, as evidenced by the fact that absolute localization judgments are less accurate and more biased for touches that are far from the nipple and the perceived location is pulled systematically toward the nipple.

### The poor spatial precision of the breast

The poor spatial precision of the breast – about four times lower than that of the hand and slightly worse than the back – replicates previous findings (9) achieved using less reliable methods (two point threshold and a ‘same-different’ paradigm). Tactile spatial precision is determined by density of innervation: more neural tissue is devoted to more highly innervated body regions, and this increased central representation is a key contributor to the increased precision. The low spatial precision of the nipple and areola is consistent with a histological study revealing these regions to be sparsely innervated (14).

### Larger breasts confer lower precision

We found a significant relationship between breast size and spatial precision: women with larger breasts tended to exhibit lower spatial precision on their breasts, consistent with previous findings that tactile precision scales with body size. Indeed, the spatial precision of the hand has been shown to depend on the size of the hand, with smaller hands exhibiting better precision (6, 7). These results are consistent with the hypothesis that the number of tactile nerve fibers does not scale with body size, so fibers are more sparsely distributed on bigger bodies, leading to lower precision. Consequently, the precision of the breast is likely determined initially by torso precision and then as a result of subsequent expansion.

### Reports are biased towards the nipple

The mental representation of the body is not uniform and veridical (15). Localization tends to be more precise when stimuli are applied near anatomical points of reference that form perceptual anchor points (16). For example, the navel and spine act as anchor points along the abdomen (17, 18): touches to the navel or spine are never mistaken for touches anywhere else on the abdomen. Furthermore, touches to locations near these two anchor points are mislocalized following a bias toward these areas: touches near the navel are pulled toward the navel and touches near the spine are pulled toward the spine. Similar biases are observed on the back of the hand, where touches are mislocalized to be closer to the fingers than they actually are (19), and on the forearm, where they are pulled toward the wrist (20). Analogously, we found that precision is highest near the nipple, and perceived locations are pulled toward the nipple, suggesting that the nipple plays a pivotal role in the mental representation of the breast. It must be noted, however, that these same biases may be inadvertently influenced by our study design. As participants were annotating 3D meshes of their breasts, the cognitive importance of the nipple may have also caused them to report sensations as closer to the nipple regardless of the actual percepts. This observation motivates standard reporting methodologies for morphologically diverse body parts such as the(21).

### Implications for breast prostheses

Understanding the spatial precision of the breast is particularly important given a recent proliferation of efforts - including using autologous tissue, synthetic grafts and neuroprosthetic or bionic approaches - to restore sensation to the breast following mastectomy (22, 23). In particular, this work shows that the spatial resolution of an implantable sensor sheet should depend on the distance from the nipple. Individual sensors should be placed in each quadrant of the nipple-areolar complex while subsequent sensors should be placed radially with increasing separation (0.5 to 2 cm inter-sensor spacing) up to a 5 cm radius at which point resolution would remain constant at approximately 2 cm. These values, however, depend on the spatial resolution of the stimulation technology which can vary significantly for both peripheral nerve stimulation (24) and intracortical microstimulation (25). Moreover, little is understood about the tactile coding of the primary afferents of the breast and further research will be needed to both inform stimulus patterning and inform safe stimulation parameters (26–28).

### Conclusion

The breast has unique sensory properties because (1) it mediates lactation and nursing, (2) it comprises distinct regions – the nipple, areola, and outer breast – which differ in the type of skin and patterns of sensory innervation, (3) it has erectile function and gives rise to erogenous sensations, and (4) it undergoes variable expansion across individuals during puberty. First, we find that spatial precision on the breast is lower than that of the back, previously regarded as the body region with lowest tactile precision. Second, spatial precision is lower in larger breasts, presumably due to sensory innervation being fixed prior to expansion during puberty. Third, the nipple itself has poor precision and yet plays a major role in how participants perceive tactile events on their breast.

## Methods

### Participants

A total of 48 healthy adult women (24.2 ± 3.1, 19-32 years) participated in this study: Thirty-four in the location discrimination task (mean ± standard deviation, age range; 22.6 ± 2.7, 19-28 years), ten in the medial breast quadrant localization task (25.3 ± 3.8, 19-32 years), and ten in the absolute localization task (24.7 ± 2.9, 20-28 years). Some participants performed multiple tasks. Experimental procedures were performed in accordance with the relevant guidelines and regulations and were approved by the Institutional Review Board of the University of Chicago (IRB 18-0135). The informed consent process included documentation of written consent from each participant, and the participants were compensated for their participation. We excluded any women who, by self-report, were currently pregnant or breastfeeding, had a history of breast surgery, or any diagnosis of neurological illness.

### Breast measurements

In addition to self-reported bra size, we collected standardized, objective measurements from each participant. Specifically, we measured the bust – i.e. the circumference of the chest at the level of the nipple – and subtracted from it the under-bust, the circumference at the level of the inframammary fold (mean ± standard deviation, range; 12.4 ± 4.6, 7.6-25.4 cm). We also measured the diameter of the areola and nipple (areola: 37.4 ± 11.9, 21-80 mm; nipple: 12.7 ± 2.7, 7-20 mm).

### Tactile Stimuli

Touches were delivered manually with a 1-mm tipped XP-Pen (XP-Pen USA, CA, USA) which allowed the experimenter to lightly press the stimulus against the skin and ensure that the skin was indented uniformly across touches. The XP-Pen was selected instead of monofilaments because, in pilot experiments, several participants reported discomfort with the sharp edges of the monofilaments on their breasts while also allowing for the application of a constant force. The participants were asked to report any discomfort with the application of the stimulus or the inability to feel the application of the stimulus reliably. Note that spatial precision is consistent across stimulus amplitudes as long as the touch is sufficiently above threshold (29), which was the case here.

### Psychophysical tasks

#### Location discrimination

Precision was tested at each of four body regions: the lateral breast, the medial breast (which includes the NAC), the thenar eminence of the hand, and the upper back. The participant – wearing a gown that exposed only the location to be tested – lay supine on a massage table for testing on the lateral breast, medial breast, and thenar eminence, and prone for the testing on the back. Unless the participant expressed a preference, the side of the body to be tested (left/right) was chosen randomly (through a random number generator) but each participant was tested on the same side for all regions. The order in which the different regions were tested was randomized to eliminate any effects of fatigue or learning.

On each trial, two touches were applied to nearby locations and the participant’s task was to indicate whether the second touch was above or below the first by pressing one of two buttons on a keypad. The location of one of the two touches (the reference) was consistent across each experimental block and the location of the second (the comparison) varied from trial to trial (each at a pre-specified distance). Fifteen comparison locations, aligned along an axis parallel to the body’s axis, were tested on the back, lateral breast, and medial breast. Touch locations were drawn on the body to ensure repeatable presentation and comparisons were placed at locations 0, 2.5, 5, 7.5, 10, 15, 20 or 40 mm above or below the reference (**Figure 1A**). Each comparison was repeated five times per block over three blocks (for a total of 15 repeats per comparison, with 225 total trials per body region) in a randomized order. Thirteen comparison locations were tested on the hand, 0, 1, 3, 6, 7.5, and 10 mm away from the reference in both directions along an axis in line with the thumb For the hand, a judgment of “above” indicated that the comparison was displaced toward the thumb relative to the reference. Each location was repeated 5 times across three experimental blocks (for a total of 15 repeats per comparison, with 195 total trials).

Performance was gauged by the proportion of times the participant judged a comparison as above the reference as a function of distance from the reference, where negative distances indicate comparison locations below the reference. Psychometric functions were fit to the data and used to compute the just noticeable difference.

#### Quadrant discrimination

The objective of this experiment was to assess the degree to which women can distinguish touches to different parts of their NAC. On each trial, the participant was touched at one of four locations on the nipple or areola (organized in a quadrant) and verbally identified the touch location using a number from 1 to 4. Each location was touched ten times per block (40 trials per block, with 80 total trials per breast region). For the areola, each touch was located halfway between the edge of the areola and the nipple. For the nipple, each touch was delivered at the edge of the nipple, in line with each point on the areola (**Figure 2A**). The testing side was chosen randomly for each participant (6 left/4 right).

#### Absolute localization

The objective of this experiment was to gauge the accuracy with which women could report where on their breast or back a touch was delivered. On the breast, touches were arranged such that the nipple was the central point and other touches radiated outwards from it (**Figure 3A**). Importantly, the two center-most touches occurred on the nipple itself. On the back, touches were arranged the same way around a central point located on the shoulder blade (**Figure 3B**). First, the participant’s skin was marked, then a three-dimensional scan of the breast or back was obtained using the EM3D application (Brawny Lads Software, LLC.) and .obj files were uploaded to Blender. The surface of the breast was then masked with a uniform layer of grey to obscure the markings. A laptop (13” screen) was positioned such that the participant could use the trackpad to interact with the 3D image (zoom, rotate, drag object) to report where the touch was experienced on the 3D rendering of their breast. Notably, in both cases landmarks were observable on the 3D models (e.g. nipple, shoulder, scapula, vertebral line). Each of the 25 stimulus locations was presented in a pseudorandomized order while the participants closed their eyes. After the touch, the participant opened their eyes and marked the perceived location of the touch on the 3D rendering of their breast or back. The participant was encouraged to manipulate the 3D model freely to obtain the best view of their breast or back on each trial. The participant moved the cursor and clicked to indicate where they perceived the touch. Reported locations accumulated on the 3D model throughout the block (during which each location was touched once.) to encourage placement of markers in accurate relative positions but were removed at the start of each of three blocks (yielding a total of 3 repeats for each of the 25 locations).

### Data Analysis

#### Spatial Precision

To quantify the spatial precision at each test location, we fit a psychometric function (see EQ1) to the probability of judging the second touch as above the first versus the relative location of the two stimuli. From these functions, we estimated the distance from the reference point required to reliably discriminate the sensation (75% performance, see EQ2).

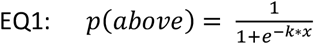

Where *k* is the growth term of the exponential function and *x* is the distance between the comparison point and the reference point.

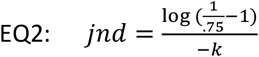

Where *k* is the growth term computed in *EQ1*.

JNDs were then compared between regions using a 1-way ANOVA followed by a Tukey’s honest significant difference post-hoc test.

#### Quadrant localization

To determine if each participant individually performed the task at greater than chance levels as well as the group as a whole, a Monte-Carlo simulation was used. Across 10,000 simulations the appropriate number of trials were simulated in which each participant randomly guessed the location and the percent correct was computed across trials for each simulation. Significance was then computed within participant by comparing the observed performance against the simulated distribution and then applying Bonferroni correction. To then determine if the whole group outperformed chance, across each of the 10,000 simulations, the cross-participant average percent correct was computed and the observed percentage compared against the resultant value. The effect size was computed by comparing the observed average percent correct with the mean of the simulation divided by the standard deviation of the simulation.

#### Absolute localization task error and bias

To compute the error for the task, we considered the 3D position of both the stimuli and response. First, to compute the total error, we computed the 3D Euclidean distance between the stimulus location and the participants response on each trial and then averaged these values together. While the skin is not flat, over the relatively short distances between stimulus and response (typically <5 cm), Euclidean distance was sufficient as minimal curvature occurred over this range. To measure the bias for each point, the 3D position for all responses for a given point across blocks was averaged together and the Euclidean distance between the resultant point and the stimulus location was averaged (centroid). If responses were randomly distributed around the stimulus location, then the error of the average response would be minimal. If there was a consistent offset, then the bias error would be similar to the total error. Finally, to compute imprecision, we computed the mean Euclidean error between the centroid and individual responses.

Linear mixed models were used to assess significant effects using Matlab’s *fitlme* function. The formula used for each test can be found in Tables 1-4. While the fixed effects varied, participants were treated as random effects and the intercept and slope were allowed to vary for each.

#### Absolute localization task angular error

To compute the angle of the errors for each response we measured the four-quadrant inverse tangent (EQ3) between each point and response. Importantly, this was only considered in 2-dimensions as the variance of angles in the 3^rd^ dimension (depth) were drastically dependent on breast size and the location of the point on the breast. Thus, the angle between the stimulus location and each response (as well as the centroid across responses) was computed. Then, for each stimulus the angle between the stimulus location and the reference point was computed (center angle) the difference in angle between said angle and the centroid angle was measured.

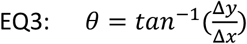

#### Absolute localization task uniformity test

To test if the cross-participant biases were uniform or not we measured the range-normalized mean squared error (rnMSE, EQ4) between the average distribution of error angles across participants and the mean of the distribution, similar to measurements of residual variance. Then, to determine if the observed rnMSE was significant, we used a permutation test in which for each permutation (n = 1e5) we shuffled the angles for each participant before averaging across participants and computing the rnMSE to produce a null distribution of rnMSEs that might be expected if the participants responses were not systematic.

Finally, we computed the proportion of simulations that had greater rnMSEs than the observed rnMSE to produce the p value which was considered significant if it was less than 0.05.

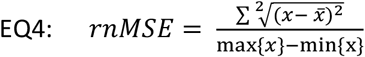

#### Absolute localization task vector strength

The vector strength for individual participants was computed using EQ5. Importantly, given the observed heteroskedasticity between distance and error, we did not incorporate the length of the vectors in the computation as this would have biased the computation towards points distal to the nipple for the breast but would not have influenced the back measurements. As the angular errors were distributed between -π and π, to test if the observed vector strength was significant, we used a Monte-Carlo simulation in which for each simulation (n = 1e5) we sampled angles from the range [-π, π] from a uniform distribution (n = 11) and computed the vector strength from each simulation. We then compared the observed vector strengths for each participant and location against the null distribution and considered it significant if the value exceeded the 95^th^ percentile of the null distribution.

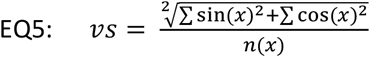

## Acknowledgments

Research supported in this publication was supported in part by National Cancer Institute grants R21CA226726 and R01CA281301, National Institute of Neurological Diseases and Stroke grant NS122333, and funding from the University of Chicago Women’s Board. The content is solely the responsibility of the authors and does not necessarily represent the official views of the National Institutes of Health. We would like to thank Katrina Schmitt and Elizabeth Pinkerton for their help on this project. SJB passed away during the writing of the manuscript.

## Data availability

Data for this study can be found online at FigShare (10.6084/m9.figshare.27939606) while code is available on GitHub (https://github.com/sensorimotor-bionics/BreastPrecision). Participant meshes are not available due to the sensitive nature of the data except for one participant who explicitly consented to a cropped version of their model being used for figures.

## Author Contributions

S.J.B and S.T.L conceived the project. C.M.G and K.H.L designed the experiments and assembled the experimental apparatus. K.H.L, E.I.BW, E.E.F, and A.F. collected the data. C.M.G performed the analyses and wrote the manuscript. All authors assisted with editing the manuscript.

## Competing Interests

SJB, KHL, and STL have patents pending based on this work.

## Supplementary figures

**Supplementary Figure 1:**
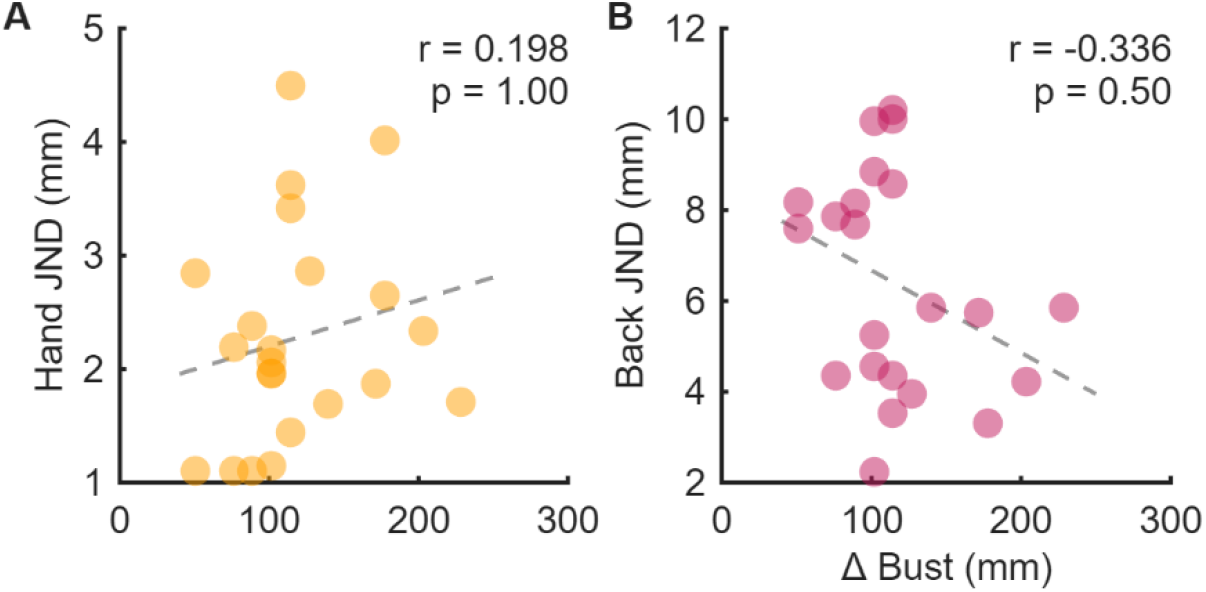
Spatial precision across regions. Relationship between the delta bust (bust – underbust) and the spatial precision (JND) for the (**A**) hand and (**B**) back.

**Supplementary Figure 2:**
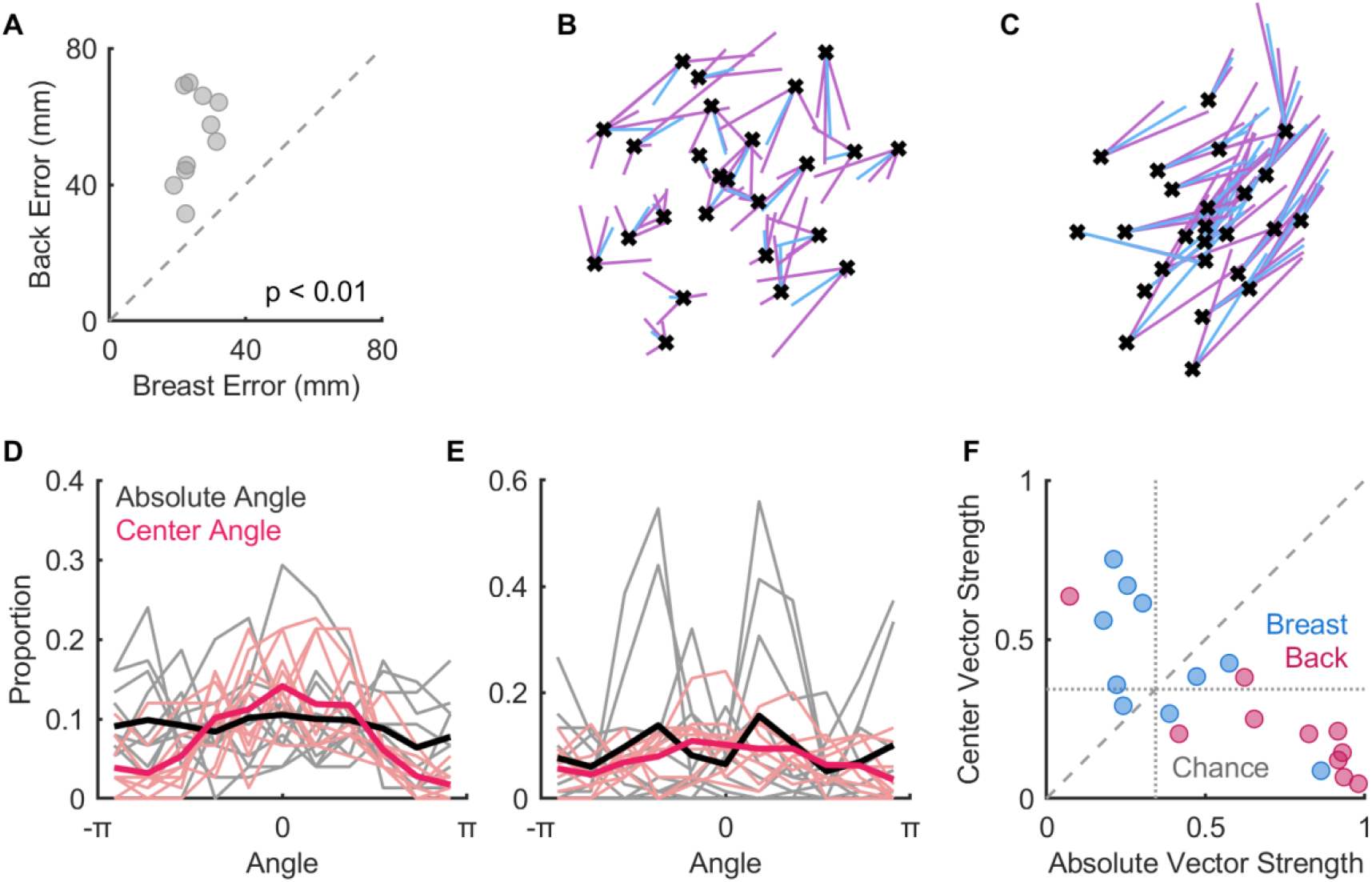
Tactile localization performance of individual participants. **A**| Mean error for each participant across distances for the back and breast. Dashed line indicates unity. **B**| Example 2D responses to localization task for the breast of one participant and the **C**| back of another. Black crosses indicate the true location of each stimulus, purple lines indicate the vector between the stimulus location and reported location, and blue lines indicate the vector between the stimulus location and average reported location across blocks. **D**| Cartesian coordinate plot of **Figure 3E** (breast) where individual participants are represented by lighter colors. **E**| same as **D** but for **Figure 3F** (back). **F**| Unimodal vector strength of individual participants biases for breast and back. Dashed line indicates unity, dotted line indicates 95^th^ percentile of vector strength from a simulated uniform distribution matched for the number of angles over which vector strength was computed after discretization (N = 11).

## Notes

https://doi.org/10.6084/m9.figshare.27939606

### Summary of Updates

Updated statistical analyses and addition of extra discussion points addressing limitations of the paper.

## References

1. F. McGlone, J. Wessberg, H. Olausson, Discriminative and Affective Touch: Sensing and Feeling. Neuron 82, 737–755 (2014).

2. G. Corniani, H. P. Saal, Tactile innervation densities across the whole body. Journal of Neurophysiology 124, 1229–1240 (2020).

3. J. C. Craig, K. B. Lyle, A comparison of tactile spatial sensitivity on the palm and fingerpad. Perception & Psychophysics 63, 337–347 (2001).

4. R. S. Johansson, A. B. Vallbo, Tactile sensibility in the human hand: relative and absolute densities of four types of mechanoreceptive units in glabrous skin. J Physiol 286, 283–300 (1979).

5. L. R. Edmondson, A. Jiménez Rodríguez, H. P. Saal, Expansion and contraction of resource allocation in sensory bottlenecks. Elife 11, e70777 (2022).

6. R. M. Peters, E. Hackeman, D. Goldreich, Diminutive Digits Discern Delicate Details: Fingertip Size and the Sex Difference in Tactile Spatial Acuity. J Neurosci 29, 15756–15761 (2009).

7. M. Wong, R. M. Peters, D. Goldreich, A Physical Constraint on Perceptual Learning: Tactile Spatial Acuity Improves with Training to a Limit Set by Finger Size. J. Neurosci. 33, 9345–9352 (2013).

8. F. Mancini, et al., Whole-body mapping of spatial acuity for pain and touch. Annals of Neurology 75, 917–924 (2014).

9. S. Weinstein, Intensive and extensive aspects of tactile sensitivity as a function of body part, sex and laterality. The skin senses (1968).

10. B. Longo, A. Campanale, F. Santanelli di Pompeo, Nipple–areola complex cutaneous sensitivity: A systematic approach to classification and breast volume. Journal of Plastic, Reconstructive & Aesthetic Surgery 67, 1630–1636 (2014).

11. G. V. Tairych, et al., Normal Cutaneous Sensibility of the Breast. Plastic and Reconstructive Surgery 102, 701 (1998).

12. J. C. Craig, K. O. Johnson, The Two-Point Threshold: Not a Measure of Tactile Spatial Resolution. Curr Dir Psychol Sci 9, 29–32 (2000).

13. A. Javed, A. Lteif, Development of the Human Breast. Semin Plast Surg 27, 5–12 (2013).

14. M. Gutiérrez-Villanueva, et al., The sensory innervation of the human nipple. Ann Anat 229, 151456 (2020).

15. M. R. Longo, Distortion of mental body representations. Trends in Cognitive Sciences 26, 241–254 (2022).

16. E. H. Weber, De Pulsu, resorptione, auditu et tactu: Annotationes anatomicae et physiologicae … (C.F. Koehler, 1834).

17. R. W. Cholewiak, J. C. Brill, A. Schwab, Vibrotactile localization on the abdomen: Effects of place and space. Perception & Psychophysics 66, 970–987 (2004).

18. J. B. Van Erp, Presenting directions with a vibrotactile torso display. Ergonomics 48, 302–313 (2005).

19. F. Mancini, M. R. Longo, G. D. Iannetti, P. Haggard, A supramodal representation of the body surface. Neuropsychologia 49, 1194–1201 (2011).

20. X. Fuchs, D. U. Wulff, T. Heed, Online sensory feedback during active search improves tactile localization. Journal of Experimental Psychology: Human Perception and Performance 46, 697–715 (2020).

21. M. Nielsen, et al., Sensory Survey 3D: an open source utility for the annotation of projected fields for sensory neural interfaces. [Preprint] (2026). Available at: https://www.researchsquare.com/article/rs-9252777/v1 [Accessed 2 April 2026].

22. S. T. Lindau, S. J. Bensmaia, Using Bionics to Restore Sensation to Reconstructed Breasts. Front. Neurorobot. 14 (2020).

23. E. H. Courtiss, R. M. Goldwyn, BREAST SENSATION BEFORE AND AFTER PLASTIC SURGERY. Plastic and Reconstructive Surgery 58, 1 (1976).

24. H. Charkhkar, et al., High-density peripheral nerve cuffs restore natural sensation to individuals with lower-limb amputations. Journal of neural engineering 15, 056002 (2018).

25. C. M. Greenspon, et al., Evoking stable and precise tactile sensations via multi-electrode intracortical microstimulation of the somatosensory cortex. Nat. Biomed. Eng 1–17 (2024). 10.1038/s41551-024-01299-z.

26. E. L. Graczyk, et al., The neural basis of perceived intensity in natural and artificial touch. Science Translational Medicine 8, 362ra142–362ra142 (2016).

27. T. G. Hobbs, et al., Biomimetic stimulation patterns drive natural artificial touch percepts using intracortical microstimulation in humans. J. Neural Eng. 22, 036014 (2025).

28. C. M. Greenspon, et al., Intracortical microstimulation in humans: a decade of safety and efficacy. [Preprint] (2025). Available at: https://www.medrxiv.org/content/10.1101/2025.08.11.25332271v1 [Accessed 15 August 2025].

29. G. O. Gibson, J. C. Craig, Relative roles of spatial and intensive cues in the discrimination of spatial tactile stimuli. Perception & Psychophysics 64, 1095–1107 (2002).

